# Immunogenicity of a new gorilla adenovirus vaccine candidate for COVID-19

**DOI:** 10.1101/2020.10.22.349951

**Authors:** Stefania Capone, Angelo Raggioli, Michela Gentile, Simone Battella, Armin Lahm, Andrea Sommella, Alessandra Maria Contino, Richard A. Urbanowicz, Romina Scala, Federica Barra, Adriano Leuzzi, Eleonora Lilli, Giuseppina Miselli, Alessia Noto, Maria Ferraiuolo, Francesco Talotta, Theocharis Tsoleridis, Concetta Castilletti, Giulia Matusali, Francesca Colavita, Daniele Lapa, Silvia Meschi, Maria Capobianchi, Marco Soriani, Antonella Folgori, Jonathan K. Ball, Stefano Colloca, Alessandra Vitelli

**Author notes:** These authors contributed equally (Stefania Capone, Angelo Raggioli).

## Abstract

The COVID-19 pandemic caused by the emergent SARS-CoV-2 coronavirus threatens global public health and there is an urgent need to develop safe and effective vaccines. Here we report the generation and the preclinical evaluation of a novel replication-defective gorilla adenovirus-vectored vaccine encoding the pre-fusion stabilized Spike (S) protein of SARS-CoV2. We show that our vaccine candidate, GRAd- COV2, is highly immunogenic both in mice and macaques, eliciting both functional antibodies which neutralize SARS-CoV-2 infection and block Spike protein binding to the ACE2 receptor, and a robust, Th1- dominated cellular response in the periphery and in the lung. We show here that the pre-fusion stabilized Spike antigen is superior to the wild type in inducing ACE2-interfering, SARS-CoV2 neutralizing antibodies. To face the unprecedented need for vaccine manufacturing at massive scale, different GRAd genome deletions were compared to select the vector backbone showing the highest productivity in stirred tank bioreactors. This preliminary dataset identified GRAd-COV2 as a potential COVID-19 vaccine candidate, supporting the translation of GRAd-COV2 vaccine in a currently ongoing Phase I clinical trial (NCT04528641).

## INTRODUCTION

Severe Acute Respiratory Syndrome CoronaVirus-2 (SARS-CoV-2) emerged in December 2019, and is responsible for the COVID-19 pandemic which has so far caused worldwide 38,202,956 confirmed cases, and 1,087,069 deaths, as of October 15^th^ 2020. Despite a growing number of both observational studies and randomized clinical trials currently running in multiple countries, to date no approved drug is licensed for the treatment or prevention of the disease, and vaccination is still regarded as a critical global priority. SARS-CoV-2 is the third novel betacoronavirus in the last 20 years to cause substantial human disease; however, unlike its predecessors SARS-CoV and MERS-CoV, SARS-CoV-2 transmits efficiently from person-to-person (1). To address the urgent need for a medical countermeasure to prevent the further dissemination of SARS-CoV-2 we have employed a novel replication defective simian adenoviral vector generated from a gorilla group C isolate: GRAd32. Viral vectored vaccines are amenable to accelerated developmental timelines due to the ability to quickly insert a foreign antigen into the E1 region of the viral genome and rescue the novel vaccine vector in an appropriate production cell line to have it ready for preclinical testing. In addition, robust GMP production processes are already available, which are easily scalable to meet the demand for a pandemic vaccine.

Simian adenoviruses derived from Chimpanzee, Bonobo and Gorilla, which are not known to infect or cause pathological illness in humans, consequently have low/no seroprevalence (0%-18%) in the human population (2). Prior studies in thousands of human subjects using simian adenoviral vaccine vectors encoding different antigens (relevant to Ebola, malaria, Hepatitis-C, HIV, and RSV), have shown that this vaccine platform is safe and can generate potent, durable, and high-quality T-cell and antibody responses (3) (4) (5) (6) (7). Furthermore, the current pandemic is giving a strong boost to the clinical validation of the simian adeno-vectored vaccine technology, since the most advanced vaccine candidate in Phase 3 clinical trial, AZD1222, is based on a chimpanzee adenoviral vector, ChAdOx1 (8).

In this study, we generated different genome deletions in the GRAd32 vector backbone and inserted a transgene cassette containing either the SARS-CoV-2 full length Spike protein (S) or its stabilized form (S-2P) (9) to preserve the neutralization-sensitive epitopes at the apex of the pre-fusion structure and to improve expression levels from transduced cells. The selected COVID-19 candidate vaccine vector, GRAd-COV2, has a genome deleted of the entire E1 and E3 regions and the native E4 region replaced with the E4 orf6 of human Adenovirus 5. We show here that this modification of the backbone improved vector productivity using a scalable process in stirred tank bioreactors which is a relevant feature in the context of a pandemic.

In this study we demonstrate that administering a single dose of GRAd-COV2 to nonhuman primates and mice we stimulate a Th1 dominant T-cell response, hACE2 receptor blocking antibodies and SARS-CoV-2 neutralizing antibodies. These results support the ongoing clinical development of GRAd-COV2 vaccine for prevention of COVID-90 19 (NCT04528641).

## RESULTS

### GRAd32 has low seroprevalence in humans

GRAd32 is a novel simian adenovirus isolated from a captive gorilla. Based on a phylogenetic analysis of aligned adenoviral polymerase sequences, GRAd32 falls into the group C adenoviruses, as the human Ad5 (Fig 1A). The prevalence of pre-existing immunity to GRAd32 vector and to the common human Ad5 was estimated in a small set of 40 human sera of US origin by a viral neutralization assay based on secreted alkaline phosphatase (SEAP) reporter gene activity (10). Greater than 32% of the serum samples were negative (IC50 titer <18) for neutralization of GRAd32. In addition, the IC50 titers of the positive sera were low and only 10% had titers greater than 200 (Fig 1B). Titers less than 200 may not be inhibitory in vivo (11). In comparison, 84% of serum samples were positive for neutralization of Ad5, with 67,5% of the positive samples having endpoint titers greater than 200 (Fig 1B).

**Fig. 1.**
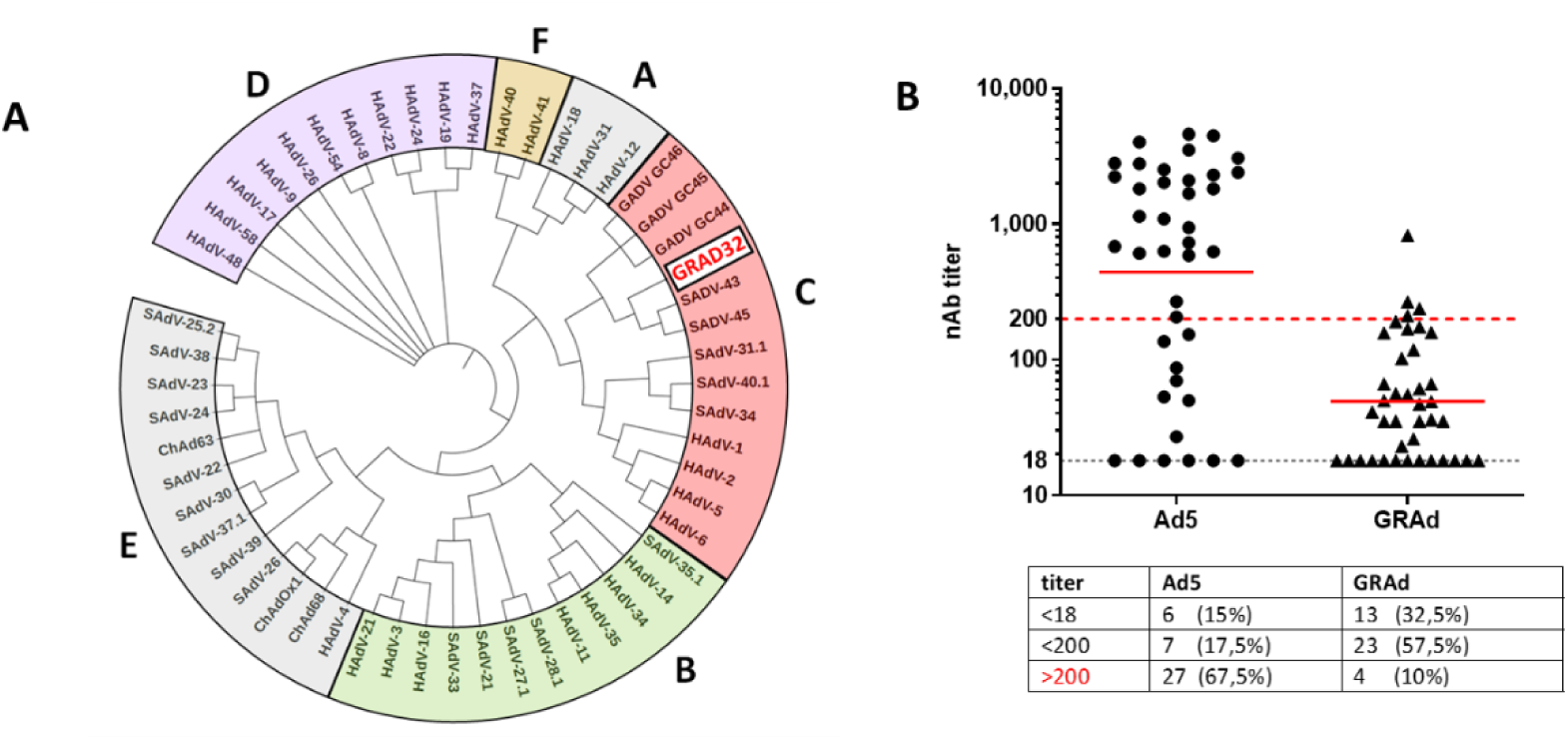
Phylogenetic analysis of GRAd32 and seroprevalence in human sera. A. Phylogenetic analysis using adenoviral polymerase sequences identifies GRAd32 as a Group C adenovirus. HAdV = Human Adenovirus, SAdV = Simian Adenovirus, GAdV = Gorilla Adenovirus B. Neutralizing antibody titers measured in sera collected from a cohort of 40 human healthy donors. Data are expressed as the reciprocal of serum dilution resulting in 50% inhibition of SEAP activity. Horizontal black dotted line indicates assay cut-off (titre of 18). Red dotted line indicates Nab titer of 200, which is reported to potentially impact on vaccine immunogenicity. Red continuous lines indicate geometric mean. The table shows the absolute numbers and the percentage of sera with NAb titers to Ad5 or GRAd32 below cut off (<18), between 18 and 200 (<200) and above 200 (>200).

### *In vitro* characterization of GRAd32-based COVID-19 vaccine constructs

We explored different GRAd32 vector backbones encoding the wild type Spike protein (S) (GenBank: QHD43416.1) or the pre-fusion stabilized S (S-2P) (9) to find the optimal combination in terms of vaccine immunogenicity and productivity. To this aim, four vaccine variants were generated, named GRAd32b-S, GRAd32b-S-2P, GRAd32c-S and GRAd32c-S-2P, which differed either in the S protein (wild type or pre-fusion) or in the vector backbone (GRAd32b: ΔE1, ΔE3 or GRAd32c: ΔE1, ΔE3, ΔE4::hAd5E4orf6) (Fig S1). In all vectors the viral E1 and E3 regions were deleted to make the virus replication defective and increase the cloning capacity; the GRAd32c vector backbone was further deleted of the E4 region, which was replaced with the E4 orf6 of human Ad5 in an attempt to optimize growth rate and yield in human cell lines, as previously described (12) (13).

The four vaccine vectors were rescued in a suspension adapted HEK293 packaging cell line and *in vitro* characterized for productivity, infectivity, genome stability and potency as measured by transgene expression. To reflect the growth conditions used for large-scale manufacturing, the productivity of the GRAd32 vector backbones was assessed after infection at controlled MOI in 2 L stirred tank bioreactors, upon collection at 48hr and 72hr post infection. As shown in Fig 2, substitution of the native E4 region with the E4orf6 of hAd5, in addition to the deletion of the E3 region, led to an improved yield (3,15e+11 vp/ml vs 7,06e+10 vp/ml) of the GRAD32c vector when harvest was collected at 60-72h after infection.

**Fig. 2.**
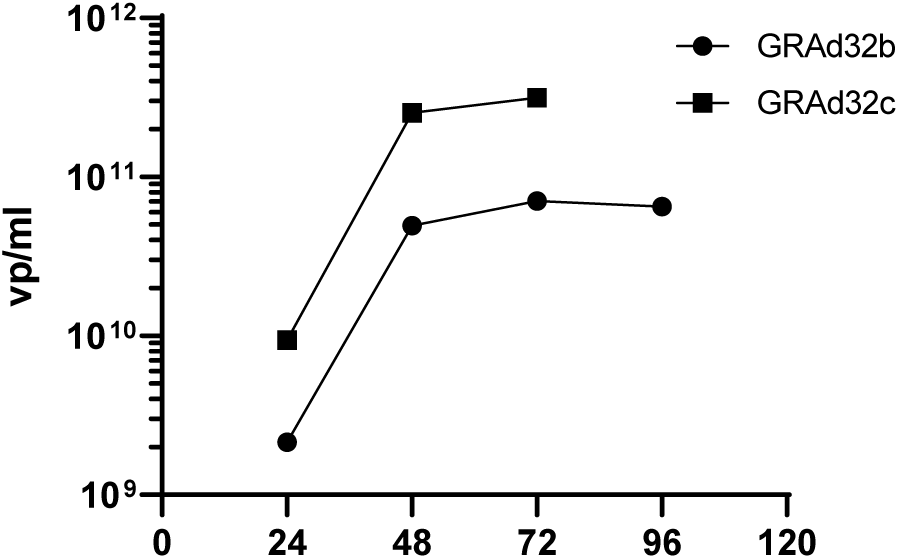
Productivity of GRAd32 backbone variants in 2L bioreactor. Suspension-adapted packaging HEK293 cells were seeded at 5e5 cells/mL and infected at MOI 200. The titer of virus contained in the bulk cell lysates collected at different time points (hours) after infection was measured by qPCR.

No difference in terms of genome stability was observed between the two vector backbones, which remained stable over 10 amplification passages as detected by restriction pattern analysis (Fig S2). However, GRAd32c showed slightly lower infectivity as measured by hexon staining in HEK293 cells respect to GRAd32b (vp/ip ratio = 178 and 63, respectively).

We then investigated the expression potency of the different vaccine vectors by HeLa cell infection and FACS analysis of whole-cell binding to either an anti-Spike S2 subunit antibody or to the soluble ACE2 receptor, and by Western Blot (WB) analysis of S protein expression in total cell lysates (Fig 3 and Fig S3). As shown in Figure 3A, 3B and 3E, FACS analysis of anti-S2 binding to whole cells revealed similar levels of cell surface-display of either S wt or S-2P, independently of the encoding GRAd32 backbone. However, in each vector backbone the displayed S-2P was clearly more abundant than the wt Spike. Such difference in the level of antigen expression was not confirmed by WB analysis of total cell lysates using an antibody directed to the receptor binding domain (RBD) or to the C terminal HA tag, possibly suggesting improved trafficking to the plasma membrane for the stabilized spike (Fig S3). Notably, a stronger binding of the recombinant soluble ACE2 receptor to the membrane-displayed S-2P protein was observed with respect to S wt (Fig 3B, 3C), suggesting that the pre-fusion conformation favors the accessibility to the RBD. Based on the above described results of *in vitro* characterization, GRAd32c-S-2P was selected as the candidate COVID-19 vaccine vector backbone given the higher vector productivity and the improved ACE2 binding of the pre-fusion stabilized Spike protein, and named GRAd-COV2.

**Fig. 3.**
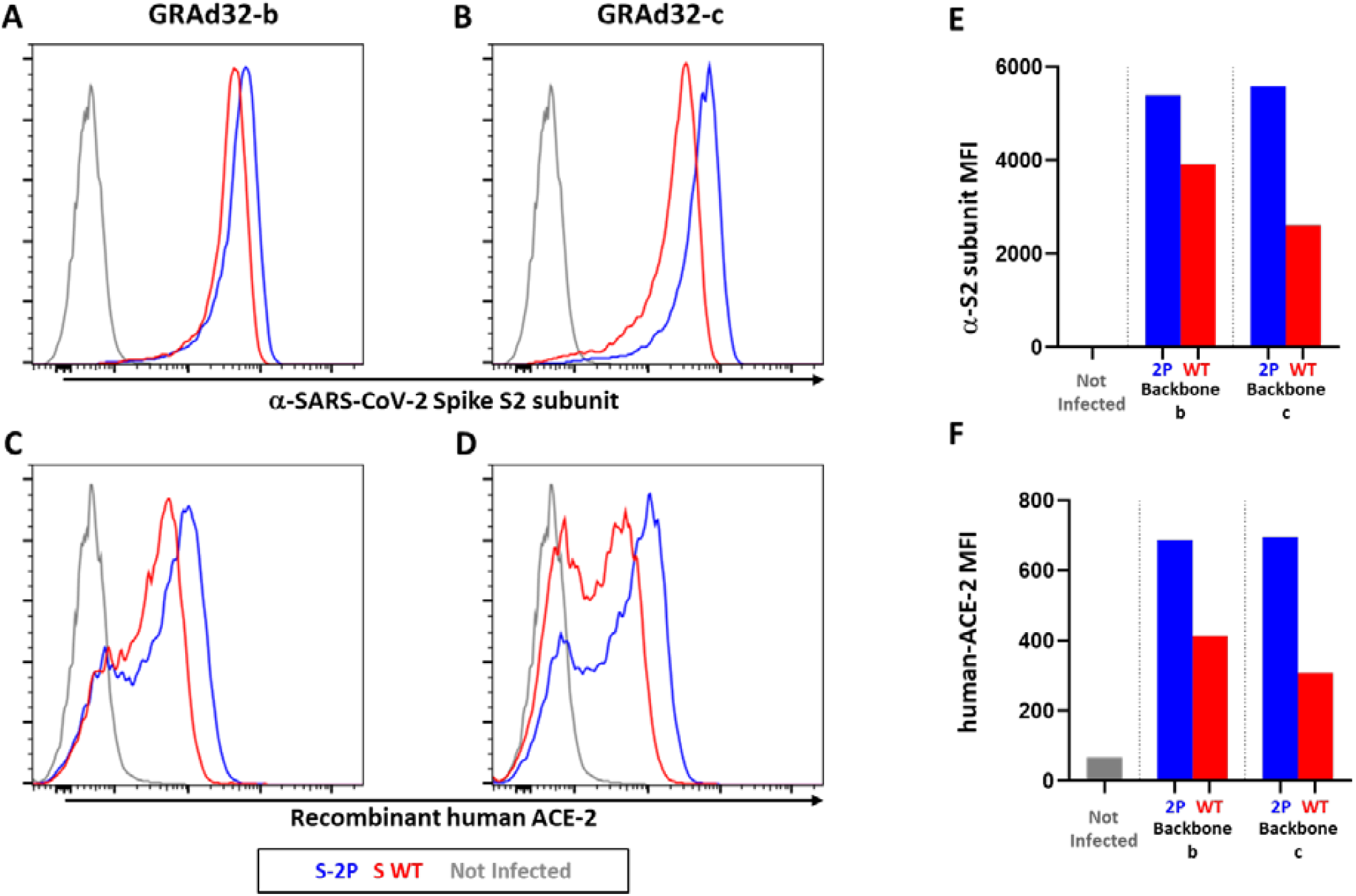
Expression of Spike antigens in GRAd32 vectors. Whole cell FACS analysis of HeLa cells not infected or infected with 150 MOI (vp/cell) of the GRAd32b (A,C) or GRAd32c (B,D). 48h after infection cells were stained with anti-S2 antibody and goat anti-mouse IgG mAb conjugated with AlexaFluor 647 (A,B) or with human soluble his-tagged ACE2 receptor and anti-His antibody conjugated with AlexaFluor 647 (C,D). The histograms of gated live cells corresponding to cells infected with the vector-encoded wt S are labeled in red and with the stabilized S-2P in blue; the grey line represents not infected cells. E, F) quantification of expression levels as detected by anti-S2 (E) or ACE-2 (F) binding, expressed as the Mean Fluorescence Intensity (MFI).

### GRAd-COV2 induces strong humoral and Th1 dominated cellular immune responses in mice

Preclinical characterization of the immunogenicity of GRAd-COV2 was performed in inbred mice. Groups of 8 BALB/c mice were vaccinated intramuscularly (IM) with a single dose of GRAd-COV2 at 1×10^9^vp. Serum samples were taken at week 2 and 5 post vaccination, and total IgG endpoint titers were measured by ELISA on full length recombinant SARS-CoV-2 Spike protein and on the Spike RBD. As shown in Fig 4A, anti-S IgG titers rose rapidly after vaccination and increased over time, and most of the antibody response was directed against the RBD. Interestingly, an anti-S IgG2a/IgG1 ratio >1 suggested a Th1 polarization of the immune responses, compared to the strongly Th2 dominated response induced by a vaccination regimen based on the recombinant S protein in alum (Fig 4B and S4).

**Fig. 4.**
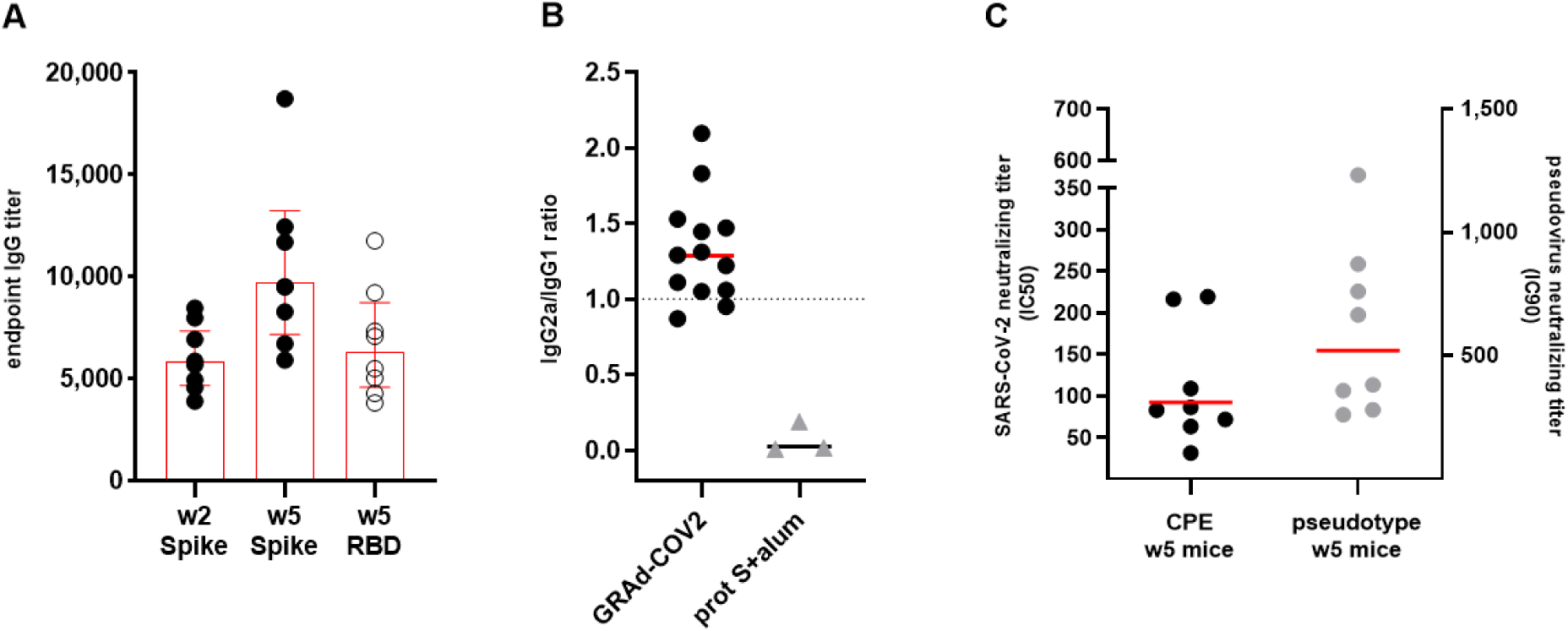
Humoral response to SARS-CoV-2 Spike induced in BALB/c mice two and five weeks after GRAd-COV2 immunization. **A**) Spike-binding total IgG titers in sera from mice immunized with 1×10^9^ vp of GRAd-COV2. IgG titers were measured in sera collected two (w2) or five weeks (w5) post immunization by ELISA on recombinant full length Spike or RBD. Data are expressed as endpoint titer. The main and error bars indicate Geometric Mean and 95% CI. **B**) The ratio between IgG2a and IgG1 titers measured in week 5 sera by ELISA on full length Spike in mice vaccinated with either 1×10^9^ vp of GRAd-COV2 (n=13) or with two injections 2 weeks apart of 2.5μg Spike protein formulated in alum adjuvant (n=3). **C**) SARS-CoV-2 neutralizing antibodies induced by GRAd-COV2 and detected in sera at week 5 post-immunization by SARS-CoV-2 (2019-nCoV/Italy-INMI) microneutralization assay on VERO E6 cells (CPE, black symbols plotted on left y axis), or by pseudotyped virus neutralization assay (grey symbols plotted on right axis). Data are expressed as IC50 or IC90, or the reciprocal of serum dilution achieving 50% or 90% neutralization respectively. Horizontal lines in panels B and C represent geometric mean.

Furthermore, the single-dose immunization with GRAd-COV2 elicited functional antibodies that neutralized the cytopathic effect (CPE) of SARS-CoV-2 on Vero E6 cells (50% blocking of CPE = 30 – 220 reciprocal serum dilution) and inhibited pseudotyped virus infection in HuH7 cell (90% inhibition of entry = 260 – 1233 reciprocal serum dilution) (Fig 4C).

Strong antigen-specific T cell responses were induced by the vaccination and measured by ELISpot and intracellular cytokine staining (ICS) from splenocytes isolated 5 weeks after immunization and stimulated with overlapping 15mer peptides. As shown in Fig 5A, IFN-γ secreting T cells recognizing epitopes in both S1 and S2 domain of Spike protein were detected (mean 1390 and 390 spot forming cells per million splenocytes respectively). In depth characterization of responding T cell subset (CD8+/CD4+) and Th polarization by ICS and FACS analysis revealed a 10% mean frequency of IFNγ secreting, Spike-specific CD8+ T cells, and a lower but clearly measurable Th1 dominated CD4+ T cell response (Fig 5B). Type 2 cytokines IL-4 and IL-13 and IL-17 secretion were also investigated by ICS and ELISpot, but were undetectable or below the limit of detection of our assays.

**Fig. 5.**
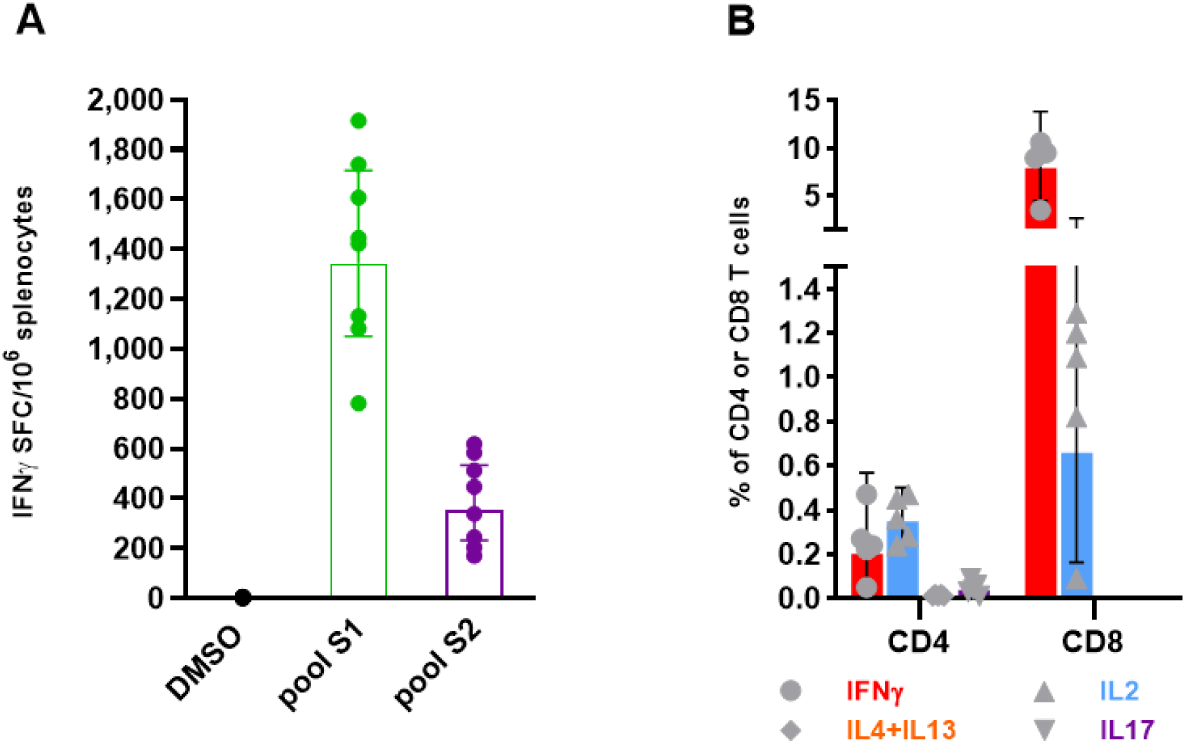
T cell response to SARS-CoV-2 Spike induced in BALB/c mice five weeks after GRAd-COV2 immunization A) IFNγ ELISpot on splenocytes. Data are expressed as IFN-γ Spot Forming Cells (SFC)/106 splenocytes. Individual data points represent response to S1 and S2 pools stimulation compared to mock stimulation (DMSO) in each animal. B) IFNγ/IL2/IL4-IL13/IL17 intracellular staining and FACS analysis on splenocytes. Data are expressed as the percentage of cytokine-secreting CD8 or CD4 T cells in response to S1 and S2 Spike peptide pools stimulation, obtained by summing reactivity to each of the 2 Spike peptide pools and subtracting 2 times the DMSO background. Main and error bars indicate Geometric mean and 95% CI in both panels A and B.

To benchmark the potency of the novel GRAd32 vector in comparison with other described simian adenoviral vectors (12), we have performed a dose-response immunization exploring a range of doses from 1×10^9^ to 1×10^5^ vp measuring IFN-γ secreting cells by ELISpot in the spleen 3 weeks after vaccination (Fig S5). Detectable immune responses in all animals were already observed at the dose 1×10^6^ vp, revealing the strong immunological potency of the new gorilla vector. Notably, strong IFN-γ secreting T cell responses were also found in the lungs of the GRAd-COV2 vaccinated animals (Fig S5).

We then compared the humoral responses induced by the wt Spike protein versus the S-2P. Sera were taken five weeks after immunization with a single administration at 1×10^9^ vp of GRAd32 vectors encoding the wt S or S-2P. We observed similar induction of total anti-S IgG in spite of significantly higher neutralization titers for the S-2P antigen (Fig 6A) measured by pseudotyped virus neutralization. Consistently, the same sera assayed for the capacity to interfere with the binding of the soluble ACE2 receptor to the immobilized RBD, showed that the S-2P antigen elicited antibodies with greater interference capacity (Fig 6B).

**Fig. 6.**
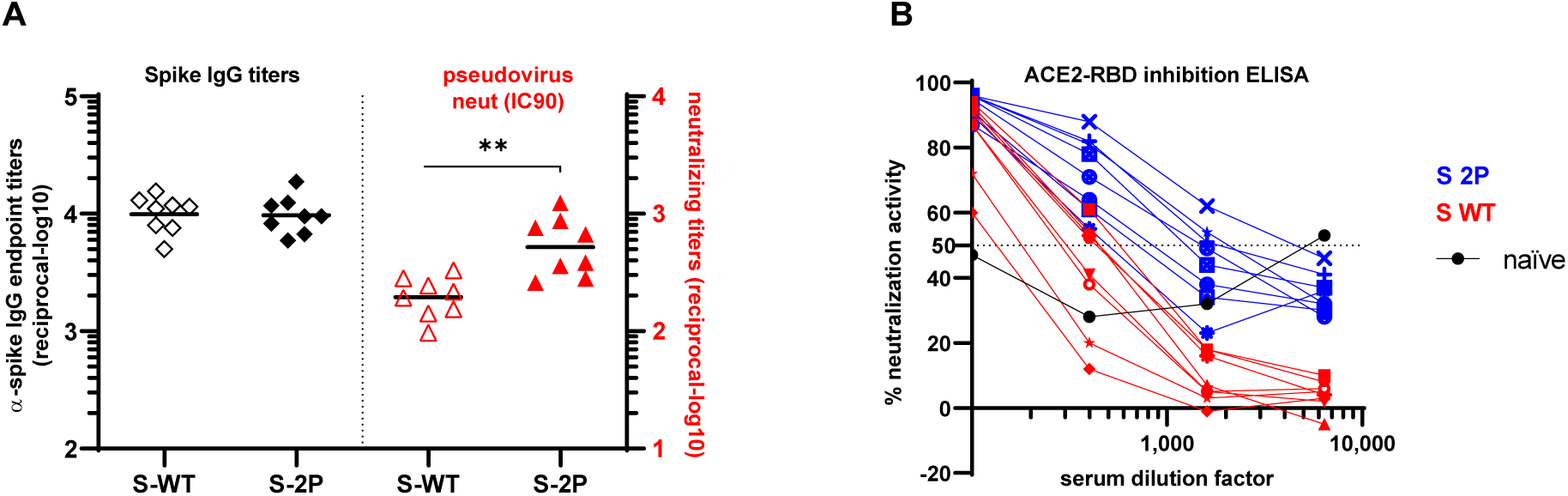
Humoral response to SARS-CoV-2 Spike induced in BALB/c mice by GRAd encoding for prefusion-stabilized versus wild type Spike A) Full length Spike-binding total IgG titers (black symbols, plotted on left y axis) and neutralizing titers on pseudotyped viruses (red symbols, plotted on right y axis) in week 5 sera from mice immunized with 1×10^9^ vp of GRAd encoding either wild type Spike (open symbols) or Spike 2P (filled symbols). Data are expressed either as endpoint titer or as IC90 neutralizing titer. Horizontal lines represent Geometric mean. Two tailed unpaired t test (** P=0.007, t=3.16, df=14), B) Inhibition of binding between immobilized RBD and recombinant human ACE2 by dilution curves of week 5 sera from mice immunized with 1×10^9^ vp of GRAd encoding either wild type Spike (red curves) or Spike 2P (blue curves). Data are expressed as percentage of neutralization activity relative to binding in the absence of serum. Inhibition curve from a naïve, non-immunized mouse as control is shown in black.

### GRAd-COV2 immunogenicity in non-human primates

Having determined that GRAd-COV2 elicits neutralizing antibodies and promotes the generation of multifunctional Th1 biased antigen-specific T cells in mice, we next evaluated the immunogenicity of the vaccine in a more relevant animal species: non-human primates. Four cynomolgus macaques were immunized intramuscularly with a single administration of 5×10^10^vp GRAd-COV2 or GRAd32c-S. Blood was drawn 1 week before vaccination and at week 2, 4, 6 and 10 after vaccination. Before vaccination all animals showed different degrees of IgG titers cross-reacting with SARS-CoV2 S and RBD, presumably deriving from previous contact with other coronaviruses (Fig 7A, 7B). However, serum IgG titers against S and RBD increased rapidly after vaccination peaking between week 2 and 4 and remained stable up to week 10 (Fig 7A, 7B). Functional antibodies measured by inhibition of SARS-CoV2 pseudotyped virus infection in Vero E6 cells and by SARS-CoV2 microneutralization (MN) assay, were also rapidly induced in all animals and peaked with different kinetics (Fig 7C, 7D). Peak Nab titers ranged between 340 - 1070 (50% CPE in MN assays) and 1580 - 4635 (IC50 pseudotyped virus entry inhibition). Importantly, vaccine-induced neutralization titers remained quite stable throughout the observation period and Nab titers measured at week 10 were comparable or higher than those measured in serum obtained from COVID-19 convalescent patients, using the same MN assay (Fig 7E).

**Fig 7.**
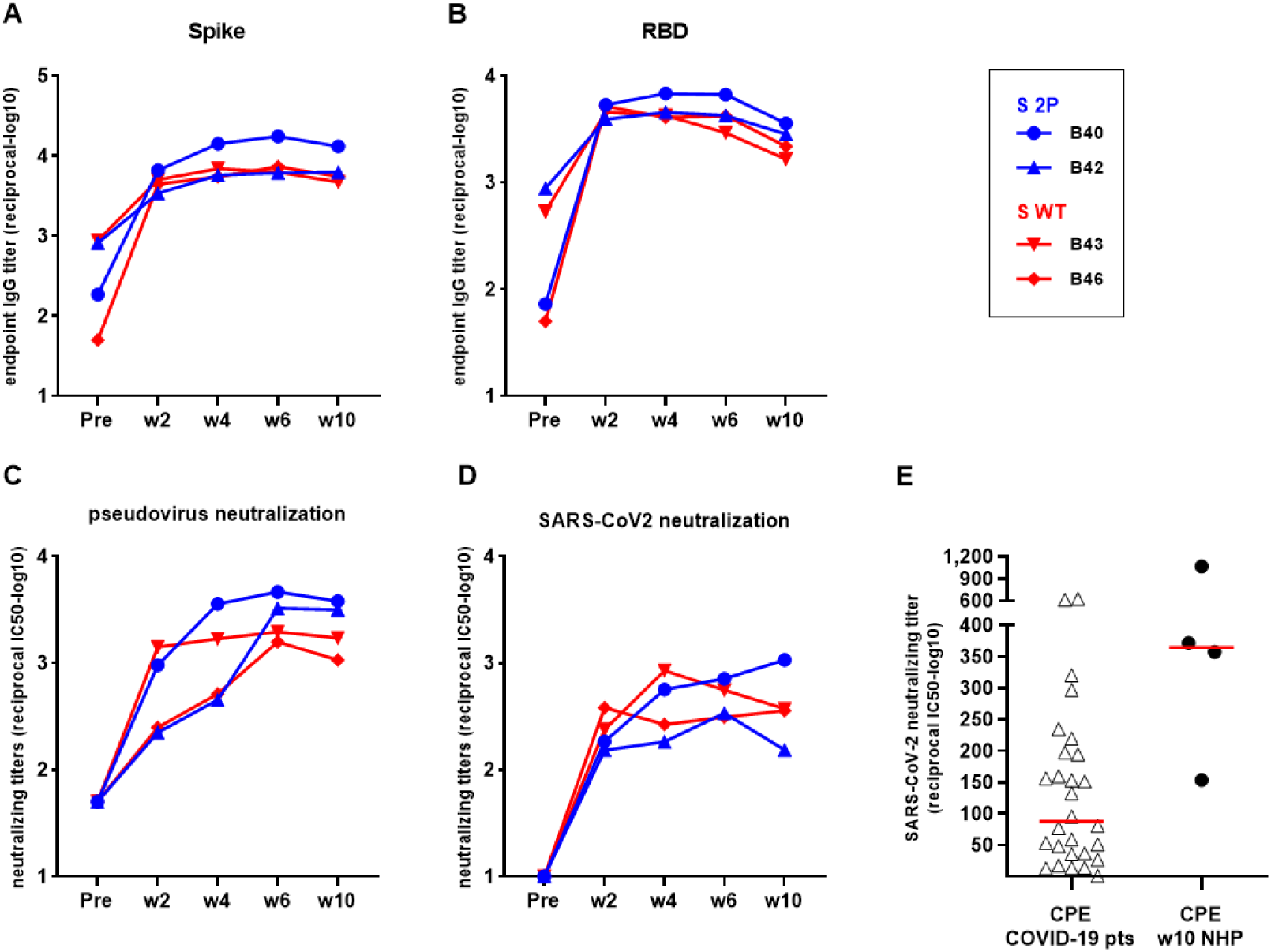
Humoral response to SARS-CoV-2 Spike induced in Cynomolgus macaques by GRAd-COV2 immunization Four cynomolgus macaques received a single 5×10^10^ vp intramuscular injection of either GRAd-COV2 (blue curves) or GRAd32c-S (red curves). Serum samples obtained before (Pre) and 2, 4, 6 and 10 weeks post-immunization were tested by A) full length Spike-binding ELISA (total IgG endpoint titers) B) RBD-binding ELISA (total IgG endpoint titers) C) pseudotyped virus neutralization assay (neutralizing IC50 titers) or D) SARS-CoV-2 microneutralization assay (neutralizing IC50 titer). E) IC50 titers measured by SARS-CoV-2 microneutralization assay in a panel of COVID-19 convalescent human sera and in GRAd-COV2 immunized macaques 10 weeks post-immunization. Horizontal lines represent geometric mean.

PBMCs were collected 2 and 8 weeks after immunization and T cell response were measured by IFNγ and IL4 ELISpot assay (Fig 8A). Strong IFN-γ secreting T cells responses were detected in all animals at week 2 (700 – 3500 SFC/10^6^ PBMCs), which somewhat contracted at week 8 but remained at high level (400 – 2500 SFC/10^6^ PBMCs). Instead, IL-4 levels were too low to be detected in macaques by ELISpot analysis (Fig 8A). The animals immunized with S-2P showed the highest number of IFN-γ secreting T cells (Fig 8B), and S1 region of the Spike protein was most immunogenic, even in this outbred animal model. Intracellular cytokine staining (ICS) performed on frozen PBMC showed that vaccination primed both CD8+ and CD4+ T cells (Fig 8C).

**Fig. 8.**
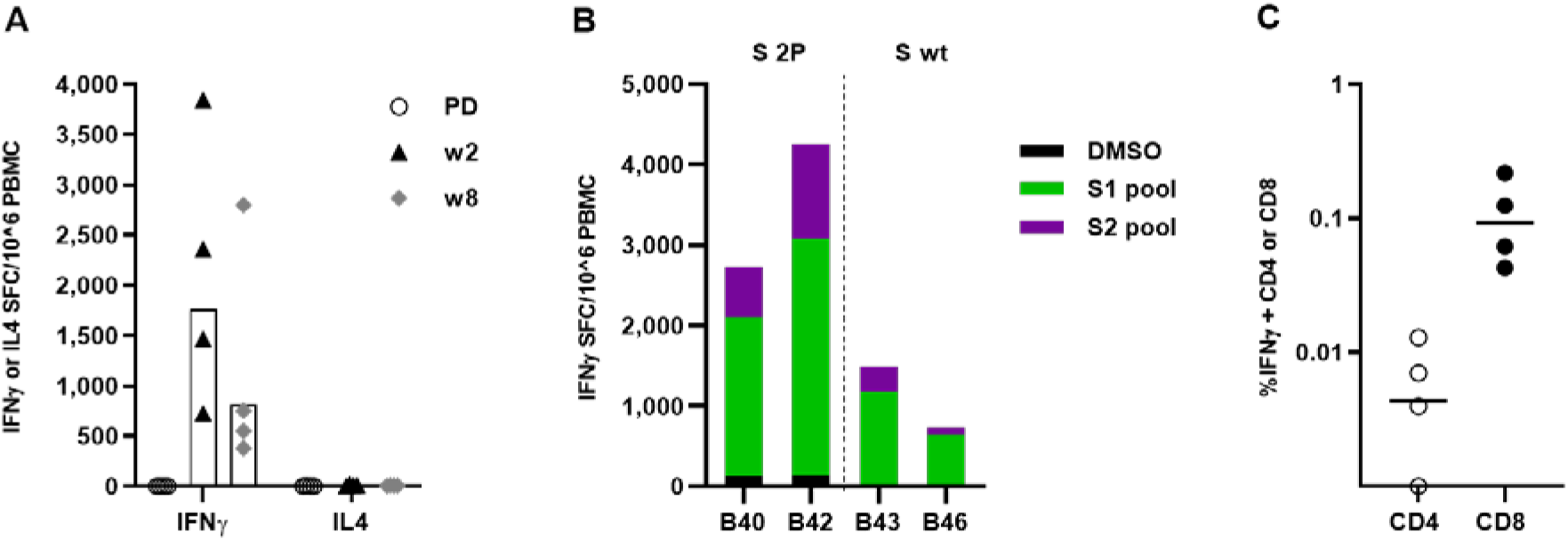
T cell response to SARS-CoV-2 Spike induced in Cynomolgus macaques upon GRAd-COV2 or GRAd32c-S immunization A) IFNγ or IL4 ELISpot on freshly isolated PBMC 2 and 8 weeks post immunization. Data are expressed as IFN-γ or IL4 Spot Forming Cells (SFC)/10^6^ PBMC. Individual data points represent cumulative Spike T cell response, calculated by summing S1 and S2 peptide pools stimulation response corrected for DMSO background of each animal. Bar represents Geometric mean. B) IFNγ ELISpot response to individual S1 and S2 peptide pools in PBMC 2 weeks post-immunization. C) intracellular staining and FACS analysis on PBMC. Data are expressed as the percentage of either CD8 or CD4 secreting cytokine in response to S1 and S2 Spike peptide pools stimulation, obtained by summing reactivity to each of the 2 Spike peptide pools and subtracting 2 times the DMSO background. Horizontal lines represent Geometric mean.

Overall, the immunogenicity of GRAd-COV2 vaccine in non-human primates is consistent with the mouse immunogenicity results and further supports the efficacy of the vaccine in promoting the generation of neutralizing antibodies and Th1 biased cellular responses.

## DISCUSSION

In this report we describe the development of a COVID-19 vaccine candidate based on a novel simian adenovirus, GRAd32, isolated from a captive gorilla and belonging to species C adenoviruses. We show here that a single immunization with the GRAd-COV2 vector encoding a pre-fusion stabilized S immunogen (S-2P) in mice and in NHP induced robust NAb responses and cellular immunity. The GRAd-COV2 encoded S-2P antigen contains the wild type leader sequence, the full-length membrane-bound S with the furin cleavage site, and two proline stabilizing mutations (9). Stabilizing the pre-fusion conformation of class I fusion proteins has successfully improved immunogenicity of these important vaccine targets for respiratory syncytial virus (14), parainfluenza virus (15), Nipah virus (16), MERS-CoV (9), and human immunodeficiency virus (17). This improvement is based on preserving neutralization-sensitive epitopes at the apex of pre-fusion structures and it was mainly shown to improve immunogenicity of secreted fusion proteins. Our data of comparison of the wt versus the stabilized full length, membrane anchored S protein extend recent preclinical studies of a vectored vaccines for SARS-CoV-2, Ad26.COV2-S, in non-human primates (18) wherein a wild type full-length S protein with proline stabilizing mutations (S.PP) represented the best antigen in terms of immunogenicity and protective efficacy in comparison with several other S antigen forms. Here we demonstrate that the GRAd vector encoding S-2P shows higher expression of membrane-anchored antigen in vitro and elicits higher titers of neutralizing antibodies in mice in comparison to the wt antigen. Similar conclusions cannot yet be drawn for non-human primates due to the very low number of animals used for this study. A more extended immunization and challenge study in NHP is ongoing to confirm these observations and, more importantly, to evaluate the protective efficacy of the vaccine. However, all the vaccinated monkeys responded rapidly to the single vaccine shot inducing a robust antibody response, which remained sustained up to 10 weeks. A longer observation period would be required to infer that the NAb responses are long lasting; however, the ongoing clinical trial of GRAd-COV2 will provide such crucial answer. A single-shot SARS-CoV-2 vaccine would have important logistic and practical advantages compared with a two-dose vaccine for mass vaccination campaigns and pandemic control. We speculate here that a species C adenoviral vector could reach this major objective based on recent reports that human Ad-5 and ChAd3 (both species C) vectors, which in mice induced the strongest T-cell responses, showed high and persistent antigen expression in the draining lymph nodes (dLNs), with low innate immunity gene activation (19). Differently, less potent vaccine vectors, like the ones based on HAdV-28 (species D), HAdV-35 (species B), and ChAd63 (species E), induced lower antigen expression associated with robust induction of innate immunity genes as determined by expression profiling in dLNs, that were primarily associated with interferon (IFN) signaling (19). However, these observations will soon receive a feedback from the clinical testing of several SARS-CoV2 vaccine candidates based on different adenoviral vectors.

Our studies do not address the safety or the possibility of vaccine-associated enhanced respiratory disease or antibody-dependent enhancement of infection (20). However, it is worth noting that the GRAd-COV2 vaccine elicited strong and Th1-biased rather than Th2-biased T cell responses in both mice and non-human primates.

Another big challenge posed by the current pandemic is the unprecedented need for scaling up vaccine manufacturing at massive scale and at record speed. To face this challenge, we generated two backbone mutants of our gorilla adenovirus: one deleted in E3 and retaining the native entire gorilla adenovirus E4 region, the other deleted in region E3 and with region E4 substituted by the E4 orf6 of human Adenovirus 5. Comparing productivity in bioreactors and stability upon many amplification passages, an optimal vector backbone (ΔE1, ΔE3, ΔE4::hAd5E4orf6) termed GRAd-COV2 has been chosen as the most promising candidate and is currently being evaluated in clinical trials.

## MATERIAL AND METHODS

### Gorilla Adenovirus Isolation, amplification and classification

Stool specimens from healthy gorilla housed in the zoo of Bristol (UK) were collected and processed for inoculation into cell cultures as previously described (12), with minor modifications. Briefly, frozen stool specimens were thawed in excess of chilled DMEM and then clarified by centrifugation followed by 0.8um and 0.22μm syringe filtration. The filtered material was inoculated into monolayers of A549 cultivated in DMEM completed with 10% Fetal Bovine Serum (FBS) and 1% Pen-Strep, and the cultures were visually monitored for cytopathic effect (CPE) for at least 28 days after inoculation. Cell monolayers showing clear sign of CPE were detached and subjected to 3 cycles of freeze/thaw (−80°C/37°C) to release the viral particles. Lysates were then clarified by centrifugation and applied onto fresh A549 to perform a large-scale preparation. Purified viral particles were obtained by Ion Exchange (VivaPure® AdenoPACKTM 20, Sartorius), and the DNA extracted and subjected to Next Generation Sequencing. Group C adenovirus assignment for GRAd32 was based on a phylogenetic analysis of aligned adenoviral polymerase sequences using a bootstrap-confirmed Maximum Likelihood tree (500 replicates) calculated with MEGA 10.1.7 (21). Tree display was performed with iTol (22). Polymerase sequences were aligned with MUSCLE (23).

### Vector construction

A pBeloBAC11-based Gorilla Adenovirus-specific shuttle BAC containing the adenoviral genome ends, a ΔE1 cassette and the adenovirus pIX gene was generated as previously described (12). Purified virus genome was isolated by Proteinase K digestion followed by phenol/chloroform extraction, and homologous recombination occurring between the right ITR and pIX present on both the shuttle BAC and the purified viral genome allowed for the insertion of the gorilla adenovirus genome in the shuttle plasmid with the deletion of the E1 region from bp 445 to 3403. This plasmid was designated as pGRAd32 ΔE1. Secreted Embryonic Alkaline Phosphatase (SEAP) coding sequence was inserted under the control of the CMV promoter (including the CMV enhancer) and the bovine Growth Hormone polyadenylation signal in the E1 locus to generate GRAd32 ΔE1 SEAP.

The pGRAd32 ΔE1 vector was further modified by recombineering to generate the ΔE1ΔE3 (named “GRAd32b”, with deletion of the E3 region from bp 28479 to bp 32001, encompassing part of the E3 12.5K, then the entire E3 CR1-alpha, E3 gp19K, E3 CR1-beta, E3 CR1-gamma, E3 RID-alpha, E3 RID-beta and E3 14.7K of the GRAd32 wild type genome) and the ΔE1ΔE3ΔE4 (named “GRAd32c”, with an additional deletion of the entire E4 region from bp 34144 to 36821, and its substitution with human Adenovirus 5 E4orf6 (12) backbones.

### Construction, amplification and purification of GRAd32 vectors expressing SARS-CoV2 spike gene

A human-codon optimized version of the SARS-CoV-2 spike coding sequence was synthesized by Doulix (Venice, Italy) and subcloned into a shuttle plasmid between the AscI and PacI restriction sites, between the tetO-hCMV promoter and the Woodchuck Hepatitis Virus (WHP) Post trascriptional Regulatory Element (WPRE) - bovine Growth Hormone poly(A) (bGHpA) sequence to generate a functional expression cassette. Two mutations were introduced to convert aa 986-987 KV into 2P, to stabilize the protein in its pre-fusion state (9). An HA tag, derived from the human influenza hemagglutinin protein, and composed of a 9-amino acid peptide sequence, Tyr-Pro-Tyr-Asp-Val-Pro-Asp-Tyr-Ala, was fused downstream of the last SARS-CoV2 S aa (Thr1273) flanked at its 5’ and 3’ side by a Gly and a Ser, respectively, to facilitate antigen expression detection by the widely commercially available HA antibodies. In addition, a minimal Kozak sequence (5’ – CCACC – 3’) was placed immediately upstream of the start codon to enable efficient initiation of translation. The cassette encoding for either SARS-CoV2 S or SARS-CoV2 S-2P was inserted by homologous recombination in the E1 locus of the GRAd32b and GRAd32c vectors, thus generating the GRAd32b-S, GRAd32b-S-2P, GRAd32c-S and GRAd32c-S-2P vectors.

All cloning PCR amplifications were performed using the Q5® High-Fidelity DNA Polymerase (New England Biolabs, Ipswich MA) according to standard procedures.

GRAd32 preAd plasmids were digested with PmeI to release the viral ITRs, and transfected in a suspension-adapted HEK293-based cell line to rescue the virus vector. Vectors were then expanded up to a production in 2L Bioreactor (Biostat B DCU, Sartorius Stedim Biotech), whereby suspension cells were seeded at 5e5 cells/mL, grown in CD293 complemented with L-Glutamine 6mM (Gibco) and Pluronic F68 0.05% (Gibco) and infected at MOI 200. The titer of virus contained in the bulk P2 cell lysates collected at 60-72h hours after the infection was measured by qPCR. Productivity was defined as the ratio between the number of viral particles quantified by qPCR in the P2 bulk lysate over the number of cells counted at the time of infection. Finally, P2 purification was performed by VivaPure® AdenoPACKTM 20 (Sartorius), and purified viral particles used for the infectivity assay (see below).

### Infectivity assay

2×10^5^ HEK293 were seeded onto 24well plates previously pre-coated with poly-L-lysine (Sigma), and infected with 2×10^5^, 1×10^5^ or 0.5×10^5^ viral particles. Infected cells were blocked at the indicated time points with ice-cold methanol, incubated with Anti-Hexon primary antibody (Abcam) followed by detection with secondary anti-Mouse IgG Peroxidase (Sigma) and Vector NovaRED substrate kit for peroxidase (Vectorlabs). The infectious unit ratio was determined as the ratio between the number of stained cells over the number of physical particles used in the assay.

### Western Blot

1×10^6^ HeLa cell, cultivated in DMEM completed with 10% fetal bovine serum (FBS) + 1% Pen/Strep were infected at MOI 150, collected at the indicated time points, and cell pellets were lysed in appropriate amount of RIPA buffer (Sigma) completed with protease inhibitor (Roche) for 30’ on ice. Cell lysates were cleared by centrifugation for 30’ at 15000rpm 4°C, and the supernatant quantified by Bradford (Biorad). 50ug of clarified lysates were loaded onto 4-12% acrylamide gel (Thermo Fisher), transferred onto iBlot 2 NC Mini Stacks membranes (Invitrogen) and incubated with the anti-HA (Abcam) or anti-RBD (Sino Biological) primary antibodies. Detection of the SARS CoV-2 spike protein was achieved by incubation with an anti-rabbit (Sigma) secondary antibody, followed by ECL (Invitrogen) incubation, and images were taken using a Chemidoc (Biorad).

### Neutralizing Antibody Assay

Neutralizing antibody (NAb) titers in human sera were assayed as previously described (10). Briefly, 8×10^4^ HEK293 cells per well were seeded in 96 well plates the day before the assay. Each adenoviral vector (Ad5 or GRAd 32) encoding for secreted alkaline phosphatase (SEAP) was preincubated for 1h at 37°C alone or with serial dilutions of control or test serum samples and then added to the 80-90% confluent 293 cells. After incubation for 1h at 37°C supernatant was then removed and replaced with 10% FBS in DMEM. SEAP expression was measured 24 hours later with the chemiluminescent substrate from the Phospha-Light™ kit (Applied Biosystems). Neutralization titers were defined as the dilution at which a 50% reduction of SEAP activity from serum sample was observed relative to SEAP activity from virus alone.

### HeLa cells infection and S2/ACE2 staining

HeLa cells were plated at 5×10^5^ cells/well in 6-well plates two days prior infection to allow adhesion, then infected with GRAd vectors at multiplicity of infection (MOI) of 150 for 48h. Cells were then harvested by pipetting with cold PBS+5mM EDTA and aliquoted in 5ml polystyrene FACS tubes (BD biosciences). Cells were then stained with Live/Dead fixable dye (Invitrogen); recombinant human His-tagged ACE-2 protein (RayBiotech) was incubated on cells before adding any antibodies, then detected with anti-His antibody (Sigma). S protein expression was detected with anti-SARS-CoV-2 S protein S2 subunit mouse mAb unconjugated (GeneTex). Antibody binding was detected with goat anti-mouse IgG mAb conjugated with AlexaFluor 647 fluorochrome (Sigma). Sample acquisition was performed on a LSR Fortessa-X20 cytofluorimeter (BD biosciences) and sample analysis was performed with FlowJo (TreeStar).

### Animals and *in vivo* procedures

Animal husbandry and all experimental procedures were approved by the local animal ethics council and were performed in accordance with national and international laws and policies on the protection of animals used for scientific purposes (UE Directive 2010/63/UE; Italian Legislative Decree 26/2014). Mouse and macaque studies were conducted under Research Projects authorized by the Italian Ministry of Health (Authorization nr. 1065/2015-PR-mouse and 984/2015-PR-macaques)

#### Mice

Six-week-old female BALB/c mice purchased from Envigo were acclimatized and housed in individually vented cages at the Plaisant animal facility (Castel Romano, Rome, Italy). Animal handling procedures (immunization and bleed) were performed under anesthesia. Animals were divided into experimental groups of 6 or 8 mice each and immunized with a single administration of different doses of GRAd-CoV2 (from 1×10^9^ vp to 1×10^5^ vp) or with two administrations two weeks apart of 2.5μg Spike protein (2019-nCoV Spike Protein-S1+S2 ECD, His tag, SinoBiological) formulated in final 20mg/ml Imject™ Alum Adjuvant (Thermo Scientific) via bilateral intramuscular injections in the quadriceps (50μl per side). After euthanasia, spleens and lungs were collected and processed into single cell suspensions. Splenocytes were isolated by mechanical disruption and resuspended in culture medium after red blood cells lysis by treatment with ACK buffer (Invitrogen). Lung lymphocytes were isolated with Lung Dissociation kit (Miltenyi) according to manufacturer instruction. In brief: whole lungs were placed in GentleMACS C tubes and incubated with provided enzymes for 30 minutes at 37°C on GentleMACS dissociator. Cell suspension was then passed on 70μM MACS SmartStrainer filters (Miltenyi); lymphocytes were purified by Percoll (GE Healthcare) gradient centrifugation and resuspended in culture medium. Isolated cells were then used for intracellular cytokine staining after in vitro re-stimulation and/or tested for IFNγ ELISPOT assay. Mice sera were obtained by standard 30 minutes blood coagulation at room temperature and centrifugation.

#### Macaques

The study was conducted at Aptuit srl, Verona in accordance with national legislation, under approval of the internal Aptuit Committee on Animal Research and Ethics. General procedures for animal care and housing are in accordance with the current Association for Assessment and Accreditation of Laboratory Animal Care (AAALAC) recommendations. Four male Cynomolgus monkeys (*Macaca fascicularis*) originating from Mauritius of approximately 3-4 years of age and 4 to 5 Kg of body weight at study start were housed in groups of 2 in stainless steel cages, with constant access to environmental enrichment devices within each cage. Conscious animals were vaccinated in deltoid muscle with a single dose of 5×10^10^ vp GRAd-COV2 (group 1, animal id B40/B42) and GRAd32c-S (group 2, animal id B43/B46). Bleedings were performed from femoral vein. Serum samples were harvested at w-1 (pre-dose-PD), w2, w4, w6, w10 post injection, obtained by 30 minute coagulation at room temperature and centrifugation and used for ELISA and *in vitro* neutralization assays. PBMCs were obtained from blood collected in Lithium Heparin tubes and isolated by separation on Ficoll gradient (LifeScience); PBMC were harvested at w-1 (PD), w2, and w8 and used fresh the same day for ex vivo IFNγ/IL4 ELISPOT, or frozen for intracellular cytokine staining.

### SARS-CoV-2 Spike and RBD ELISA

SARS-CoV-2 ELISA was developed in-house using His tagged proteins bound on Ni-NTA HisSorb Strips or Plates (Qiagen). ELISA assay on mouse sera were performed with His-tagged SARS-CoV-2 full length Spike protein (produced in baculovirus, SinoBiological), while ELISA with monkey sera were performed with a trimeric his-tagged stabilized SARS-CoV-2 Spike protein produced in-house in Expi293 cells or with a commercial RBD (ACROBiosystems). In brief: optimized amount of proteins in 100μl volume were bound to Ni-NTA plates/strips for 1h at 25°C in shaking (1 μg/ml for full length Spike, 2 μg/ml for R121, 0.5 μg/ml for RBD); plates were then incubated with serial dilutions of sera for 2h at 25°C in shaking, then binding was detected with specific alkaline phosphatase-conjugated secondary antibodies (anti-mouse total IgG and anti-monkey total IgG from Sigma and anti-mouse IgG1 and IgG2a from BD Biosciences). Alkaline phosphatase substrate was prepared by dissolving SigmaFast (Sigma) tablets in sterile distilled H2O. Absorbance was read at 405 and 620 nm using EnSight multiple plate reader (PerkinElmer). The endpoint titer was defined as the highest serum dilution that resulted in an absorbance value (OD-optical density) just above calculated background: for mice experiments, 3-fold the OD of 1:100 serum dilution from naïve mice; for monkey experiments, 4-fold the OD from secondary antibody alone. Serum from naïve BALB/c animals are routinely tested at 1:100 dilution as negative control in ELISA, and the resulting OD is nearly at the level of secondary Ab alone, therefore for mouse experiments baseline titers cannot be detected, calculated and plotted.

### RBD/ACE-2 neutralization ELISA

RBD/ACE-2 neutralization ELISA (ACROBiosystems) was performed according to manufacturer instruction. In brief: MaxiSorp ELISA plates (Nunc) were coated with 50ng/well (100μl at 0.5 μg/ml) of SARS-CoV-2 RBD protein for 16h at 4°C, blocked for 1.5h at 37°C, incubated 1h at 37°C with 50μl of 0.12 μg/ml biotinylated recombinant human ACE-2 protein alone or with serial dilutions of mice sera. ACE-2 binding was detected with HRP-conjugated streptavidin and developed with TMB substrate. Absorbance was read at 450nm using EnSight multiple plate reader (PerkinElmer). Data are expressed as neutralization curves calculating the percentage of ACE2/RBD binding inhibition at the different serum dilutions relative to the control without inhibitory antibodies.

### SARS-CoV2 microneutralization assay

Human sera derived from post-convalescent COVID-19 patients after signing of an informed consent, in the frame of a project aimed at following up patients.

Sera collected from patients and vaccinated animals were heat-inactivated at 56°C for 30 minutes, and titrated in duplicate in two-fold serial dilutions. Equal volumes of 100 TCID50 SARS-CoV-2 (strain 2019-nCoV/Italy-INMI) and serum dilutions, were mixed and incubated at 37 °C for 30 min. Subsequently, 96-well tissue culture plates with sub-confluent Vero E6 cell monolayers were infected with 100 μl/well of virus-serum mixtures and incubated at 37 °C and 5% CO_2_. After 48 hours, microplates were observed at the microscope for the presence of cytopathic effect (CPE). To standardize inter-assay procedures, positive control samples showing high and low neutralizing activity were used for each microneutralization assay. The supernatant of each plate was carefully discarded and 120 μl of a Crystal Violet solution containing 2% Formaldehyde was added to each well. After 30 min fixation, the fixing solution was removed and cell viability measured by photometer at 595 nm (Synergy HTX Biotek). The serum dilution inhibiting 50% of the CPE (IC50) was calculated using graph pad prism 7 as described in (24)

### SARS-CoV-2 pseudotyped virus neutralization assay

Pseudotyped viruses were produced as previously described (25). Briefly, 1.5 × 10^6^ HEK293T cells were seeded overnight in a 10 cm diameter Primaria-coated dish (Corning). Transfections were performed using 2 µg each of the murine leukemia virus (MLV) Gag-Pol packaging vector (phCMV-5349), luciferase encoding reporter plasmid (pTG126) and plasmid encoding the same human-codon optimized version of the SARS-CoV-2 spike coding sequence as detailed above, minus the HA tag. These were mixed with 24 µl cationic polymer transfection reagent (polyethylenimine), in the presence of Optimem (Gibco), and the media replaced with 10 ml complete DMEM after 6 h. A no-envelope control (empty pseudotype) was used as a negative control in all experiments. Supernatants containing SARS-CoV-2 pseudotypes were harvested at 72 h post-transfection and filtered through 0.45-μm-pore-size membranes. For infectivity and neutralization assays, either 1.5×10^4^ HuH7 cells or 2×10^4^ VeroE6 cells/well were plated in white 96-well tissue culture plates (Corning) and incubated overnight at 37°C. The following day, SARS-CoV-2 pseudotypes were incubated with heat-inactivated sera for 1 hr at RT before being added to cells for 4 h. Following this, sera and media were discarded, and 200 µl DMEM was added to the cells. After 72 h, media was discarded, and cells were lysed with cell lysis buffer (Promega) and placed on a rocker for 15μmin. Luciferase activity was then measured in relative light units (RLUs) using a FLUOstar Omega plate reader (BMG Labtech) with MARS software. Each sample was tested in triplicate.

### ELISpot

For ELISpot assays, MSIP 96 well plates were from Millipore (Multiscreen filter plates) and anti-mouse or anti-monkey IFNγ or IL4 capture and detection antibodies were all from U-CyTech. Lymphocytes (from mouse spleens/lungs or macaque PBMC) were plated in duplicate at appropriate cell densities and cultured for 18-20 hours at 37°C, 5% CO_2_ with peptide pools covering the entire SARS-CoV-2 spike protein referred to as S1 or S2 (JPT PepMix). Stimulation conditions were 1 (for mouse lymphocytes) or 3 μg/ml (for monkey’s PBMCs) final single peptide concentration, or the equivalent amount of DMSO, the peptide diluent, as negative control. Cytokine production and spot formation was detected with alkaline phosphatase-conjugated streptavidin and NBT/BCIP alkaline phosphatase substrate. The number of spots per well, directly related to the precursor frequency of antigen-specific T cells, was counted and analysed with a CTL ImmunoSpot reader. Data are expressed as cytokine spot forming cells (SFC) per million splenocytes, lung lymphocytes or PBMC, and are shown upon subtraction of DMSO background.

### Intracellular cytokine staining and FACS analysis

Mouse splenocytes were used freshly isolated, while monkey PBMCs were stimulated after thawing and overnight resting in culture medium. Cells were plated at 1×10^6^ per well in a 96-well round bottom plate and stimulated with either SARS-CoV-2 peptide pool S1 or S2, or with DMSO as negative control, all in presence of anti-CD28 antibody, for 18h (mouse cells) or 5h (monkey cells) in the presence of Brefeldin-A (Sigma). Cells were then washed to stop the stimulation, stained with Live/Dead fixable dye (Invitrogen), fixed and permeabilized with BD Cytofix/Cytoperm kit following manufacturer instructions, and then stained with specific fluorochrome conjugated antibodies. Sample acquisition was performed with LSR Fortessa-X20 cytofluorimeter (BD biosciences) and sample analysis was performed with FlowJo (TreeStar). Mouse Antibodies: CD3 AlexaFluor647, CD4 BV421, CD8 BUV395, IL-17a PerCP-Cy 5.5, IFNγ BV650, IL2 APC-R700 (from BD Biosciences), IL-4 PE, IL-13 PE (Invitrogen). NPH Antibodies: CD3 AF700, CD4 PerCP-Cy 5.5, CD8 PE (from BD Biosciences), IFNγ FITC (U-Cytech).

### Statistics

GraphPad Prism version 8 for Windows (GraphPad Software, San Diego, California, USA) was used for graphs and statistical analysis. In figure 6, since the data showed Gaussian distribution as per all normality tests run, a two-tailed unpaired Student’s t test was used. Only statistically significant results were reported in the figures, *p ≤ 0,05; **p ≤ 0,01; ***p ≤ 0,001; ****p ≤ 0,0001. For the macaque study, the numerosity of the groups was too small to allow statistical analysis.

## Acknowledgements

JKB, RAU and TS were supported by grants from the Medical Research Council UK (MR/R010307/1 and MR/S009434/1), which are both part of the EDCTP2 programme supported by the European Union, The University of Nottingham Campaign and Alumni Relations Office research donations award and the EU H2020 programme (Project 727393 –PaleBlu)

## REFERENCE

1. Tu YF, Chien CS, Yarmishyn AA, Lin YY, Luo YH, Lin YT, et al. A Review of SARS-CoV-2 and the Ongoing Clinical Trials. Int J Mol Sci. 2020;21(7).

2. Vitelli A, Folgori A, Scarselli E, Colloca S, Capone S, Nicosia A. Chimpanzee adenoviral vectors as vaccines - challenges to move the technology into the fast lane. Expert Rev Vaccines. 2017;16(12):1241–52.

3. Green CA, Scarselli E, Sande CJ, Thompson AJ, de Lara CM, Taylor KS, et al. Chimpanzee adenovirus- and MVA-vectored respiratory syncytial virus vaccine is safe and immunogenic in adults. Sci Transl Med. 2015;7(300):300ra126.

4. O’Hara GA, Duncan CJ, Ewer KJ, Collins KA, Elias SC, Halstead FD, et al. Clinical assessment of a recombinant simian adenovirus ChAd63: a potent new vaccine vector. J Infect Dis. 2012;205(5):772–81.

5. Borthwick N, Ahmed T, Ondondo B, Hayes P, Rose A, Ebrahimsa U, et al. Vaccine-elicited human T cells recognizing conserved protein regions inhibit HIV-1. Mol Ther. 2014;22(2):464–75.

6. Tapia MD, Sow SO, Lyke KE, Haidara FC, Diallo F, Doumbia M, et al. Use of ChAd3-EBO-Z Ebola virus vaccine in Malian and US adults, and boosting of Malian adults with MVA-BN-Filo: a phase 1, single-blind, randomised trial, a phase 1b, open-label and double-blind, dose-escalation trial, and a nested, randomised, double-blind, placebo-controlled trial. Lancet Infect Dis. 2016;16(1):31–42.

7. Swadling GF, Lebedev SV, Hall GN, Patankar S, Stewart NH, Smith RA, et al. Diagnosing collisions of magnetized, high energy density plasma flows using a combination of collective Thomson scattering, Faraday rotation, and interferometry (invited). Rev Sci Instrum. 2014;85(11):11E502.

8. Folegatti PM, Ewer KJ, Aley PK, Angus B, Becker S, Belij-Rammerstorfer S, et al. Safety and immunogenicity of the ChAdOx1 nCoV-19 vaccine against SARS-CoV-2: a preliminary report of a phase 1/2, single-blind, randomised controlled trial. Lancet. 2020;396(10249):467–78.

9. Pallesen J, Wang N, Corbett KS, Wrapp D, Kirchdoerfer RN, Turner HL, et al. Immunogenicity and structures of a rationally designed prefusion MERS-CoV spike antigen. Proc Natl Acad Sci U S A. 2017;114(35):E7348–E57.

10. Aste-Amezaga M, Bett AJ, Wang F, Casimiro DR, Antonello JM, Patel DK, et al. Quantitative adenovirus neutralization assays based on the secreted alkaline phosphatase reporter gene: application in epidemiologic studies and in the design of adenovector vaccines. Hum Gene Ther. 2004;15(3):293–304.

11. Buchbinder SP, Mehrotra DV, Duerr A, Fitzgerald DW, Mogg R, Li D, et al. Efficacy assessment of a cell-mediated immunity HIV-1 vaccine (the Step Study): a double-blind, randomised, placebo-controlled, test-of-concept trial. Lancet. 2008;372(9653):1881–93.

12. Colloca S, Barnes E, Folgori A, Ammendola V, Capone S, Cirillo A, et al. Vaccine vectors derived from a large collection of simian adenoviruses induce potent cellular immunity across multiple species. Sci Transl Med. 2012;4(115):115ra2.

13. Brough DE, Lizonova A, Hsu C, Kulesa VA, Kovesdi I. A gene transfer vector-cell line system for complete functional complementation of adenovirus early regions E1 and E4. J Virol. 1996;70(9):6497–501.

14. McLellan JS, Chen M, Joyce MG, Sastry M, Stewart-Jones GB, Yang Y, et al. Structure-based design of a fusion glycoprotein vaccine for respiratory syncytial virus. Science. 2013;342(6158):592–8.

15. Stewart-Jones GBE, Chuang GY, Xu K, Zhou T, Acharya P, Tsybovsky Y, et al. Structure-based design of a quadrivalent fusion glycoprotein vaccine for human parainfluenza virus types 1-4. Proc Natl Acad Sci U S A. 2018;115(48):12265–70.

16. Dang HV, Chan YP, Park YJ, Snijder J, Da Silva SC, Vu B, et al. An antibody against the F glycoprotein inhibits Nipah and Hendra virus infections. Nat Struct Mol Biol. 2019;26(10):980–7.

17. Pancera M, Zhou T, Druz A, Georgiev IS, Soto C, Gorman J, et al. Structure and immune recognition of trimeric pre-fusion HIV-1 Env. Nature. 2014;514(7523):455–61.

18. Mercado NB, Zahn R, Wegmann F, Loos C, Chandrashekar A, Yu J, et al. Single-shot Ad26 vaccine protects against SARS-CoV-2 in rhesus macaques. Nature. 2020.

19. Quinn KM, Zak DE, Costa A, Yamamoto A, Kastenmuller K, Hill BJ, et al. Antigen expression determines adenoviral vaccine potency independent of IFN and STING signaling. J Clin Invest. 2015;125(3):1129–46.

20. Graham BS. Rapid COVID-19 vaccine development. Science. 2020;368(6494):945–6.

21. Kumar S, Stecher G, Li M, Knyaz C, Tamura K. MEGA X: Molecular Evolutionary Genetics Analysis across Computing Platforms. Mol Biol Evol. 2018;35(6):1547–9.

22. Letunic I, Bork P. Interactive Tree Of Life (iTOL) v4: recent updates and new developments. Nucleic Acids Res. 2019;47(W1):W256-W9.

23. Edgar RC. MUSCLE: multiple sequence alignment with high accuracy and high throughput. Nucleic Acids Res. 2004;32(5):1792–7.

24. Ferrara F, Temperton N. Pseudotype Neutralization Assays: From Laboratory Bench to Data Analysis. Methods Protoc. 2018;1(1).

25. Tighe PJ, Urbanowicz RA, Fairclough L, McClure CP, Thomson BJ, Gomez N, et al. Potent anti-SARS-CoV-2 Antibody Responses are Associated with Better Prognosis in Hospital Inpatient COVID-19 Disease. medRxiv. 2020:2020.08.22.20176834.

