## supplementary info for "Immunogenicity of a new gorilla adenovirus vaccine candidate for COVID-19"

**Supplementary data**

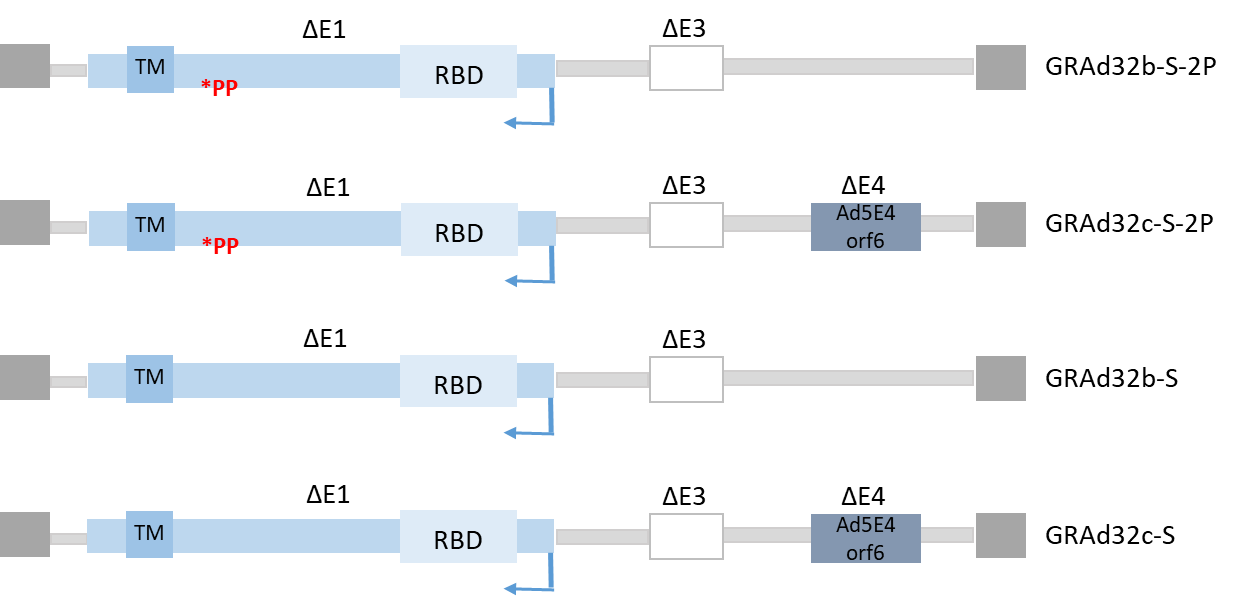

**Fig S1**. Schematic cartoon of GRAd32 genome backbones and encoded Spike antigens. The full length wild type (S) or the two proline-stabilized (S-2P) Spike gene expression cassette was cloned in E1 in the leftward orientation. The GRAd32b backbone is deleted of E1 and E3 regions, while GRAd32c genome is deleted of E1, E3 and E4 and replaced with the hAd5 E4 orf6

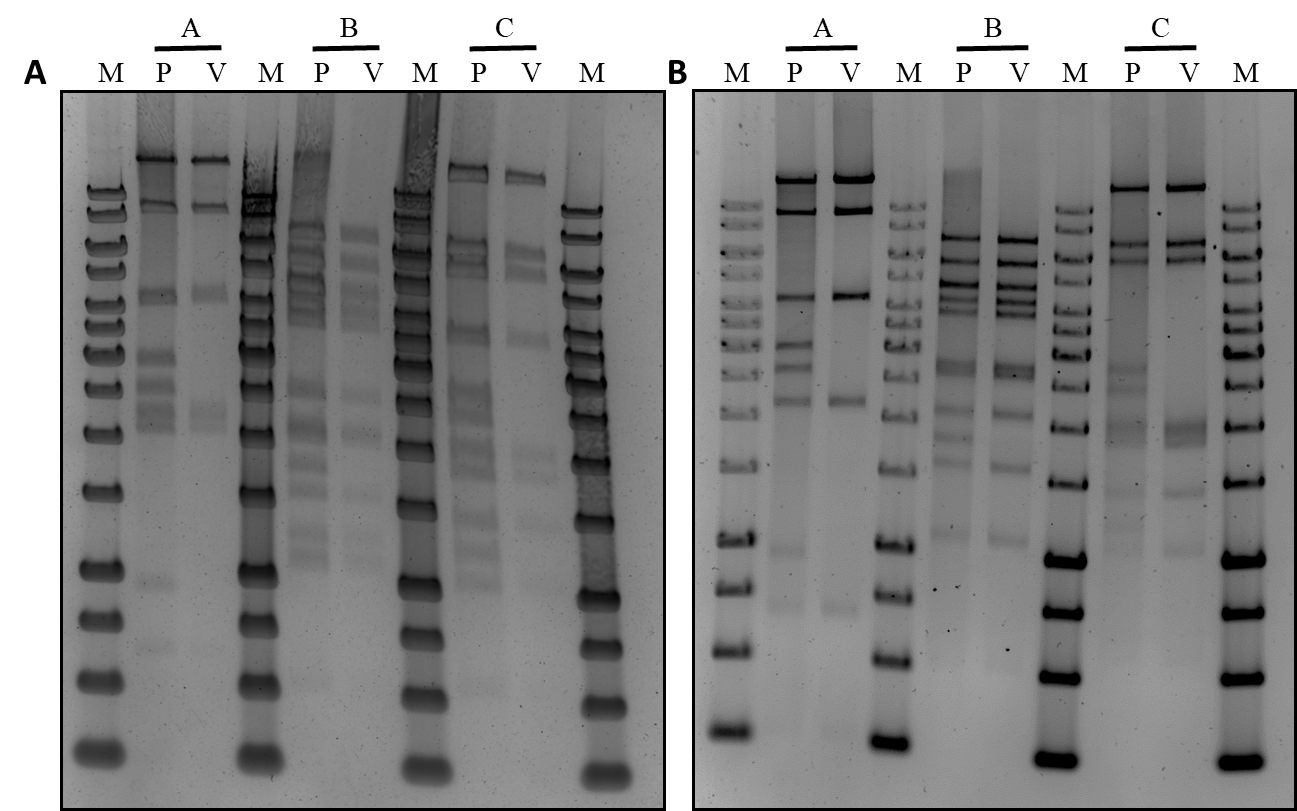

**Fig. S2** Restriction analysis of viral genome after 10 passages. Genomic DNA was extracted from GRAd32b-S-2P (A) and GRAd32c-S-2P (B) purified viral particles (V), digested with different restriction enzymes and compared to the pre-adeno plasmid (P). M: 1kb marker, BsrGI+SpeI (lane 2-3) XhoI+SphI (lane 5-6) XmnI (lane 7-8)

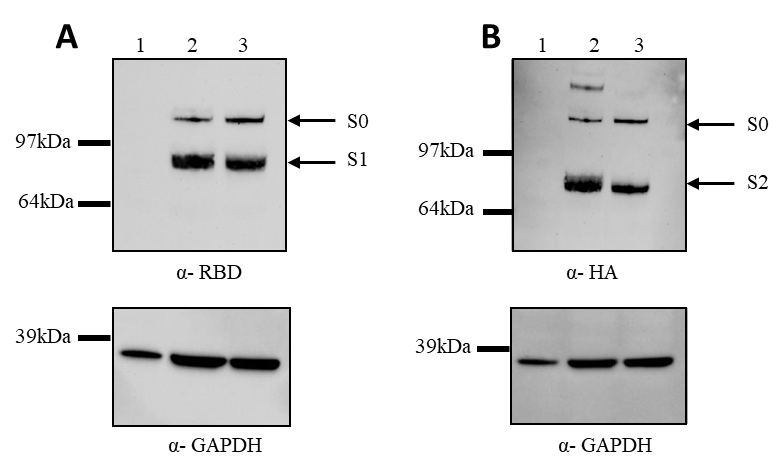

**Fig. S3** Expression of Spike antigens. Western blot analyses of reducing SDS-PAGE of cell lysates from HeLa cells not infected (lane 1) or infected with 150 MOI (vp/cell) of GRAd32 vectors encoding Spike 2P (lane 2) or Spike wt (lane 3), using either an anti-Spike RBD rabbit polyclonal antibody (A) or an anti-HA tag rabbit monoclonal antibody (B).

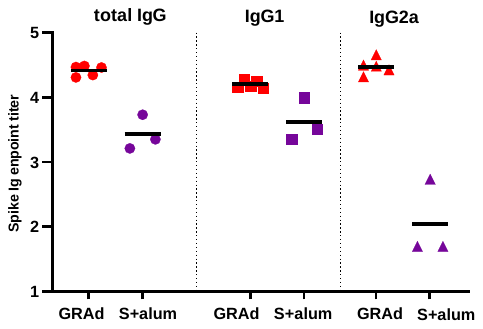

**Fig. S4** A single GRAd-COV2 administration induces similarly anti Spike IgG1 but more efficiently IgG2a than two administration of a Spike protein formulated in alum adjuvant. A) Spike-binding total IgG, IgG1 and IgG2a titers in sera from mice immunized with 1x109 vp of GRAd-COV2 or with two injections 2 weeks apart of 2.5μg Spike protein formulated in alum adjuvant. Sera collected at w4 post first immunization were tested by ELISA on recombinant full length Spike. Data are expressed as endpoint titer. Horizontal lines indicate Geometric Mean

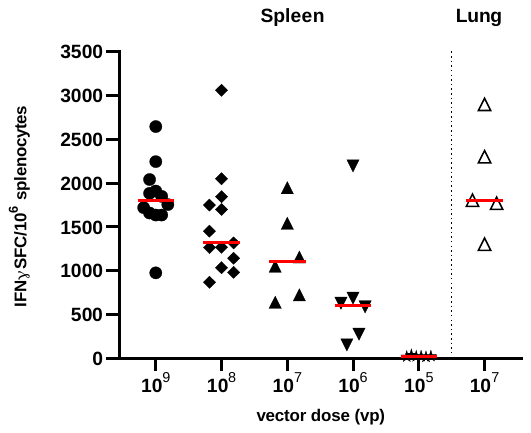

**Fig. S5** Dose-response T cell response in BALB/c mice three weeks after GRAd-COV2 vaccination. BALB/c mice received a single intramuscular injection of between 109 to 105 vp of GRAd-COV2, and spleens were collected 3 weeks post immunization. IFNγ ELISpot was done on splenocytes (filled symbols) and lung infiltrating lymphocytes (open symbols, 107 vp dose only). Data are expressed as IFN-γ Spot Forming Cells (SFC)/106 splenocytes. Individual data points represent total Spike response in each animal, obtained by summing reactivity to each of the 2 Spike peptide pools and subtracting 2 times the DMSO background. Red lines represent Geometric mean.
